# Tightly Knotted Enzymes Inhibit Protein-Protein Aggregation

**DOI:** 10.64898/2026.06.12.731949

**Authors:** Susmita Sarkar, Hemanth Mandya Nagaiah, Michael L. Klein, Vincenzo Carnevale

## Abstract

Neurodegenerative diseases are closely linked to aberrant protein aggregation arising from failures in cellular proteostasis, yet the physical determinants governing transitions between soluble states, liquid–liquid phase separation (LLPS), and aggregation remain incompletely understood. Here, we investigate how protein backbone topology influences phase behavior using ubiquitin C-terminal hydrolase L1 (UCH-L1), a highly neuron-enriched deubiquitinase in the ubiquitin–proteasome system harboring a rare, evolutionarily conserved knotted backbone topology, and its Parkinson’s disease–associated I93M mutant. Through multiscale molecular dynamics (MD) simulations of single-chain and multichain systems, we show that knot integrity acts as a conformational constraint that limits access to expanded states, and suppresses LLPS propensity. Destabilization of the native knotted ensemble in I93M reshapes the conformational ensemble, enhancing intermolecular contacts, strengthening hydrophobic interaction, and reducing solvation penalties, thereby stabilizing protein-rich phases. Within condensates, these changes lead to persistent interchain contacts, increased topological entanglement, and slower relaxation dynamics, indicative of a transition toward viscoelastic assemblies, whereas intact topology maintains dynamic, liquid-like behavior. Our results identify topological integrity as a key physical determinant of protein phase behavior and establish a mechanistic link between topological stability and condensate material properties, with implications for aggregation-associated neurodegeneration.

## Introduction

Neurodegenerative diseases, including Alzheimer’s disease (AD) and Parkinson’s disease (PD), are characterized by progressive neuronal loss and the pathological accumulation of misfolded protein aggregates[1, 2, 3]. A unifying molecular feature of these disorders is the failure of protein homeostasis, wherein the ubiquitin-proteasome system (UPS) and autophagy pathways become overwhelmed and fail to efficiently clear aberrant proteins[4]. Deubiquitinases (DUBs) play central regulatory roles within the UPS by controlling ubiquitin chain dynamics and maintaining the free monoubiquitin pool required for proteasomal substrate processing[5, 6]. Although dysregulation of DUB activity is implicated across a broad spectrum of human diseases, including neurodegeneration, cancer, and metabolic disorders[7, 8, 9], the molecular principles by which individual DUBs contribute to proteostasis failure[10, 11, 5], and how their structural properties shape aggregation susceptibility, remain incompletely understood.

Ubiquitin C-terminal hydrolase L1 (UCH-L1) is among the most neuron-enriched members of the DUB family, constituting 1–2% of total soluble brain protein with near-exclusive expression in neurons and neuroendocrine cells[12, 13, 14].Beyond its canonical role in maintaining monoubiquitin homeostasis, UCH-L1 participates in synaptic maintenance, vesicle trafficking, and long-term neuronal plasticity [15, 16, 9]. Loss of UCH-L1 activity causes axonal dystrophy and neurodegeneration in vivo[17, 18, 13], while altered expression, oxidative modification, and mutation are linked to AD and PD pathology[9, 14, 19]. The familial I93M mutation, originally identified in PD patients[20], confers a dominant toxic gain-of-function that dramatically increases aggregation propensity without significantly altering the native crystal structure[19]—highlighting a fundamental mechanistic gap: how can a nominally folded protein acquire strong aggregation propensity without large-scale structural disruption?

We propose that the answer lies in a structural feature of UCH-L1 that has received limited attention as a functional determinant: its deeply knotted 5_2_ backbone topology[21, 22, 23, 24]. UCH-L1 belongs to the rare class of proteins in which the polypeptide chain forms a non-trivial self-entanglement, with one segment threading through a loop formed by another part of the chain. Knotted proteins[25, 26, 27, 28] occur far less frequently than expected from random polymer statistics, implying strong evolutionary pressure against topological complexity[29, 30]. Nevertheless, knotting is strictly conserved across UCH-family deubiquitinases over large phylogenetic distances, implying strong positive selection for a functional advantage conferred by the knotted architecture. Previous studies of UCH-L1 have primarily focused on its enzymatic activity, substrate recognition, and disease-associated mutations[20, 7, 16, 8, 6, 31, 18, 15, 13, 9, 14, 19], while its knotted backbone topology has received comparatively limited attention as a functional determinant. The I93M mutation, however, directly challenges this view: it maps near the knotted core, and its dominant aggregation phenotype—disproportionate to large-scale structural rearrangement—points instead to a mechanism in which perturbation of backbone topology[32, 33], reshapes the conformational ensemble to promote aggregation.

In parallel, LLPS has emerged as a central organizing principle in cell biology[34, 35, 36, 37], underpinning the biogenesis of membraneless organelles and increasingly recognized as a key driver of pathological protein condensation in neurodegeneration[36, 38, 39, 40, 41, 42]. Current understanding of LLPS emphasize intrinsically disordered regions (IDRs)[43] and weak multivalent interactions[44, 45, 46, 47], casting structured proteins as passive condensate clients rather than active drivers of phase separation.

This prevailing view omits a fundamental physical parameter: protein backbone topology. From a polymer-physics standpoint, topology imposes global constraints that shape conformational entropy, modulate intramolecular contact formation, and influence intermolecular encounter geometries[48, 49]—constraints that could critically determine whether proteins remain soluble, phase-separate, or aggregate in crowded cellular environments.

Here, we address this gap by probing the relationship between backbone knot integrity, LLPS, and aggregation propensity in UCH-L1 through extensive molecular dynamics (MD) simulations of the wild-type (WT) protein and the disease-associated I93M variant (Fig. 1). By combining single-chain conformational analysis with multichain simulations across all-atom (AA) and coarse-grained (CG) resolutions, we show that backbone topology acts as an active regulator of phase behavior in a folded protein. Our results reveal a mechanistic link between topological stability and emergent collective behavior in protein assemblies, establishing backbone topology as a previously unrecognized determinant of condensate behavior and aggregation susceptibility. We introduce the concept of topological proteostasis—the maintenance of non-trivial backbone topology as a protective mechanism against aberrant phase separation and aggregation—and propose that its disruption represents a distinct failure mode in neurodegeneration.

**Figure 1:**
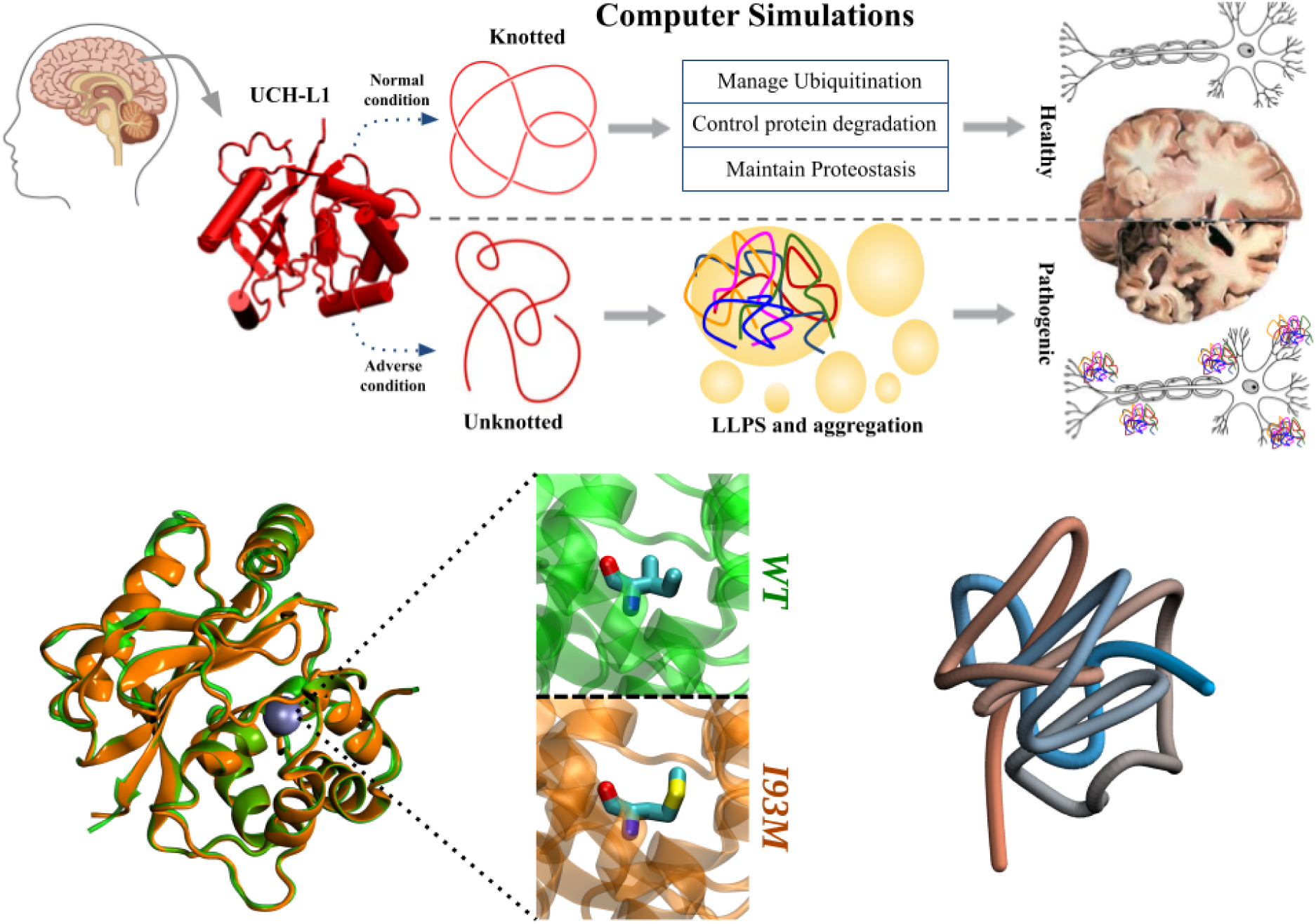
Structural and topological features of UCH-L1 and conceptual framework of this study. Ubiquitin C-terminal hydrolase L1 (UCH-L1) is a neuron-enriched deubiquitinase implicated in neurodegenerative disease and characterized by a rare 5_2_ knotted backbone topology. (A) Schematic overview illustrating the conceptual framework of this work. We propose that the knotted backbone topology stabilizes the native conformational ensemble of UCH-L1, while its perturbation by the I93M mutation shifts the conformational landscape toward interaction-prone states, promoting phase separation and altered condensate behavior that may be associated with disease conditions. Superimposed crystal structures of WT (green) and I93M mutant (orange) UCH-L1 are shown in cartoon representation, highlighting the conserved globular architecture, with the mutation site indicated as a gray sphere. A zoomed-in view of residue 93 in WT (Ile) and mutant (Met) is shown in licorice representation (C, cyan; N, blue; O, red; S, yellow), illustrating local structural differences. A smoothed backbone representation emphasizes the deeply embedded 5_2_ knot within the protein core.

## Results

### Topological regulation of UCH-L1 conformational ensembles

To determine how the I93M mutation reshapes the conformational landscape of UCH-L1, we first performed AA MD simulations of both WT and I93M monomers. Analysis of backbone root mean square deviation (RMSD) distributions relative to the corresponding crystal structures reveals a clear broadening in the mutant, with an extended high-RMSD tail indicative of enhanced sampling of expanded conformations and increased conformational heterogeneity relative to WT (Fig. 2A). CG simulations, which access longer timescales, confirm a significantly increased radius of gyration (R*_g_*) in the mutant relative to WT (Fig. 2B), alongside elevated root-mean-square fluctuations (RMSF) distributed across multiple structural regions rather than confined to the mutation site (Fig. 2C). This global redistribution indicates a weakening of intrachain stabilizing interactions rather than a locally isolated perturbation.

**Figure 2:**
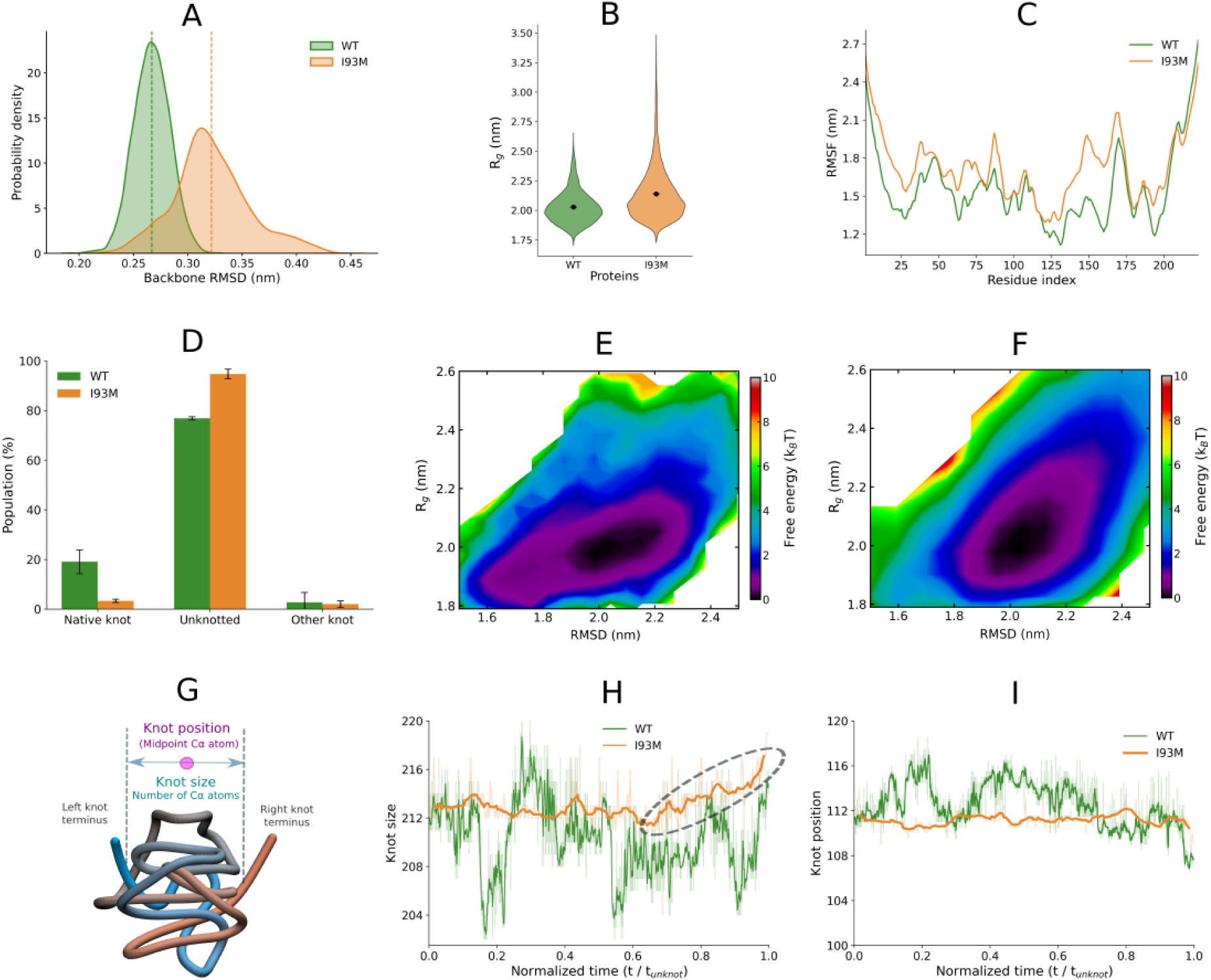
The I93M mutation destabilizes the knotted topology and reshapes conformational ensembles of mutant UCH-L1. (A) Probability distributions of backbone RMSD relative to the corresponding crystal structures from AA simulations of WT (green) and I93M (orange) UCH-L1. (B) Distribution of R*_g_* from CG simulations. (C) Residue-level flexibility measured by RMSF from CG simulations. (D) Populations of different topological states, including native knotted, alternative knot types, and unknotted configurations for WT and I93M. (E–F) Free-energy landscapes projected onto RMSD and R*_g_* for WT (E) and I93M (F). (G) Schematic representation of knot-core analysis. The knot core defines the minimal backbone segment preserving topology, with N- and C-terminal regions shown in blue and red, respectively. Knot size and position are defined from this region (see Methods). (H) Knot-size dynamics represented as a function of normalized time relative to the characteristic topological transition time (*t/t*_unknot_) for WT and I93M. (I) Knot-core positional dynamics along the backbone represented as a function of normalized time relative to the characteristic topological transition time (*t/t*_unknot_) for WT and I93M.

Given the deeply embedded 5_2_ knot of UCH-L1, we next asked whether these conformational changes are coupled to alterations in topological state. Using the Jones polynomial[50] to characterize knot status (see Methods), we found that the I93M mutation significantly reduces the fraction of fully knotted conformations, with a corresponding increase in partially and fully unknotted states (Fig. 2D). Free-energy landscapes projected onto backbone RMSD and R*_g_* (Fig. 2E-F) visualize the thermodynamic consequences: WT UCH-L1 occupies a well-defined basin at low RMSD and low R consistent with a native-like ensemble (Fig. 2E), whereas the I93M mutant populates an altered landscape in which free-energy minima shift toward higher RMSD and larger R (Fig. 2F). This population shift indicates thermodynamic stabilization of expanded, conformationally heterogeneous states rather than mere broadening around the native basin, suggesting that knot destabilisation and chain expansion are thermodynamically coupled consequences of the I93M substitution rather than independent effects.

Detailed analysis of knot dynamics reveals mechanistically distinct behaviour between the two variants (Fig. 2G–I). Profiles normalized with respect to the characteristic topological transition time (*t/t*_unknot_) show that WT UCH-L1 exhibits stable fluctuations in knot size, whereas the I93M mutant samples progressively larger knot dimensions, indicative of knot swelling (Fig. 2H). Complementary analysis of knot-core position further reveals pronounced positional fluctuations in WT, consistent with sliding motions that accommodate local thermal rearrangements (Fig. 2I). In contrast, the mutant exhibits reduced knot positional mobility despite the concurrent increase in knot size. This combination of knot swelling and reduced positional mobility is mechanistically significant: it indicates a weakening of the topological constraint and expansion of the knotted region, promoting loss of native knot integrity. Together, these results establish that the knotted backbone of WT UCH-L1 acts as an internal topological constraint that penalizes chain expansion and stabilizes the compact native ensemble, while the I93M substitution erodes this constraint to enrich a population of expanded, topologically loosened states.

### Mutation-Induced Topological Loosening Enhances Intermolecular Association

We next examined whether topological destabilization propagates to aberrant intermolecular interactions. AA simulations of preformed dimers show that WT dimers predominantly populate configurations with large center-of-mass (COM) separations, consistent with transient intermolecular association, whereas I93M dimers show a distribution strongly enriched at lower COM distances along with stable interfacial configurations (Fig. 3A,B). CG simulations initiated from spatially separated monomers confirm that the I93M mutant exhibits substantially shorter times to first association (Fig. 3C) and higher bound-state occupancy (Fig. 3D), demonstrating more rapid intermolecular encounter and more stable dimer formation.

**Figure 3:**
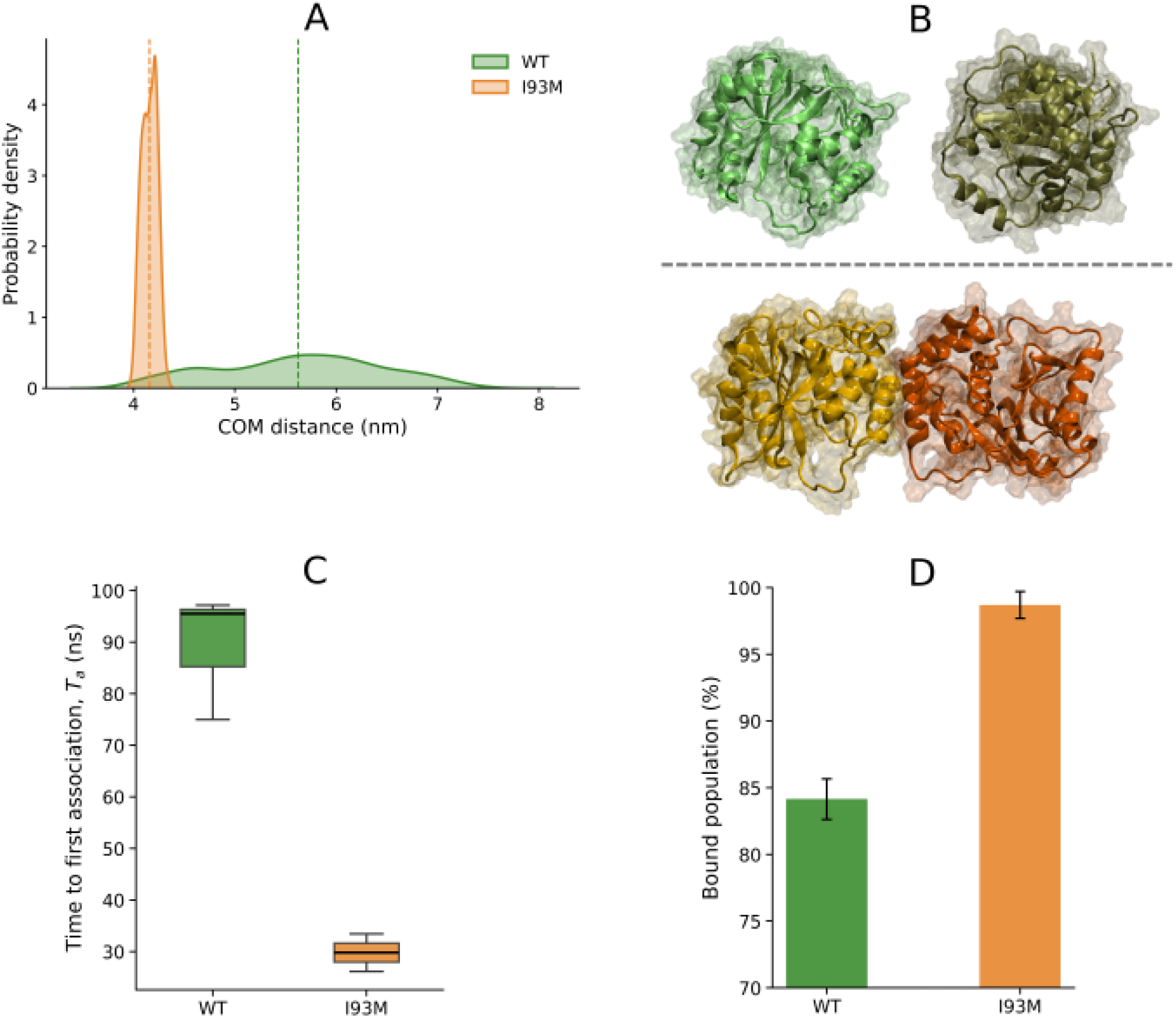
Dimerization dynamics of WT and I93M UCH-L1. (A) Probability distributions of interchain center-of-mass (COM) distance from AA simulations of preformed WT (green) and I93M (orange) dimers. (B) Representative snapshots from AA simulations of preformed WT and I93M dimers illustrating interchain orientation and interfacial stability. (C) Time to first association (*T_a_*) obtained from CG simulations initiated from spatially separated monomers, quantifying the kinetics of initial intermolecular encounter for WT and I93M. (D) Bound-state occupancy from CG simulations, quantifying the fraction of simulation time that the chains remain associated for WT and I93M.

The enhanced association is associated with a redistribution of conformational populations toward more expanded states that are enriched in the mutant and associated with enhanced intermolecular encounters. These conformations, identified in the monomer simulations, arise from reduced topological constraint and increased chain expansion, thereby increasing the likelihood of productive intermolecular encounters in the dimer simulations. In WT UCH-L1, such expanded conformations are only sparsely populated and thus contribute minimally to intermolecular association. In the I93M mutant, their statistical weight is significantly increased, amplifying the contribution of these states and shifting the balance from transient encounters to persistent dimer formation. Thus, backbone topology acts as a regulator of intermolecular affinity by controlling access to conformational sub-ensembles that govern the earliest stages of multichain assembly.

### The I93M Mutation Enhances LLPS in Multichain Systems

To investigate how molecular-level changes propagate to collective phase behavior, we performed large-scale CG simulations of multichain systems(34 protein copies; ∼500 µM) initiated from homogeneous dispersions. Both WT and mutant UCH-L1 form protein-rich condensates, but with notable quantitative and qualitative differences. Protein concentration distributions reveal an order-of-magnitude difference between dilute and dense phases for both variants—a hallmark of LLPS—but the I93M system shows substantially higher dense-phase concentration and lower dilute-phase concentration, indicating enhanced partitioning into the condensed phase (Fig. 4A). Representative simulation snapshots further supports more pronounced self-assembly and larger, more compact condensates in the I93M system (Fig. 4B-C).

**Figure 4:**
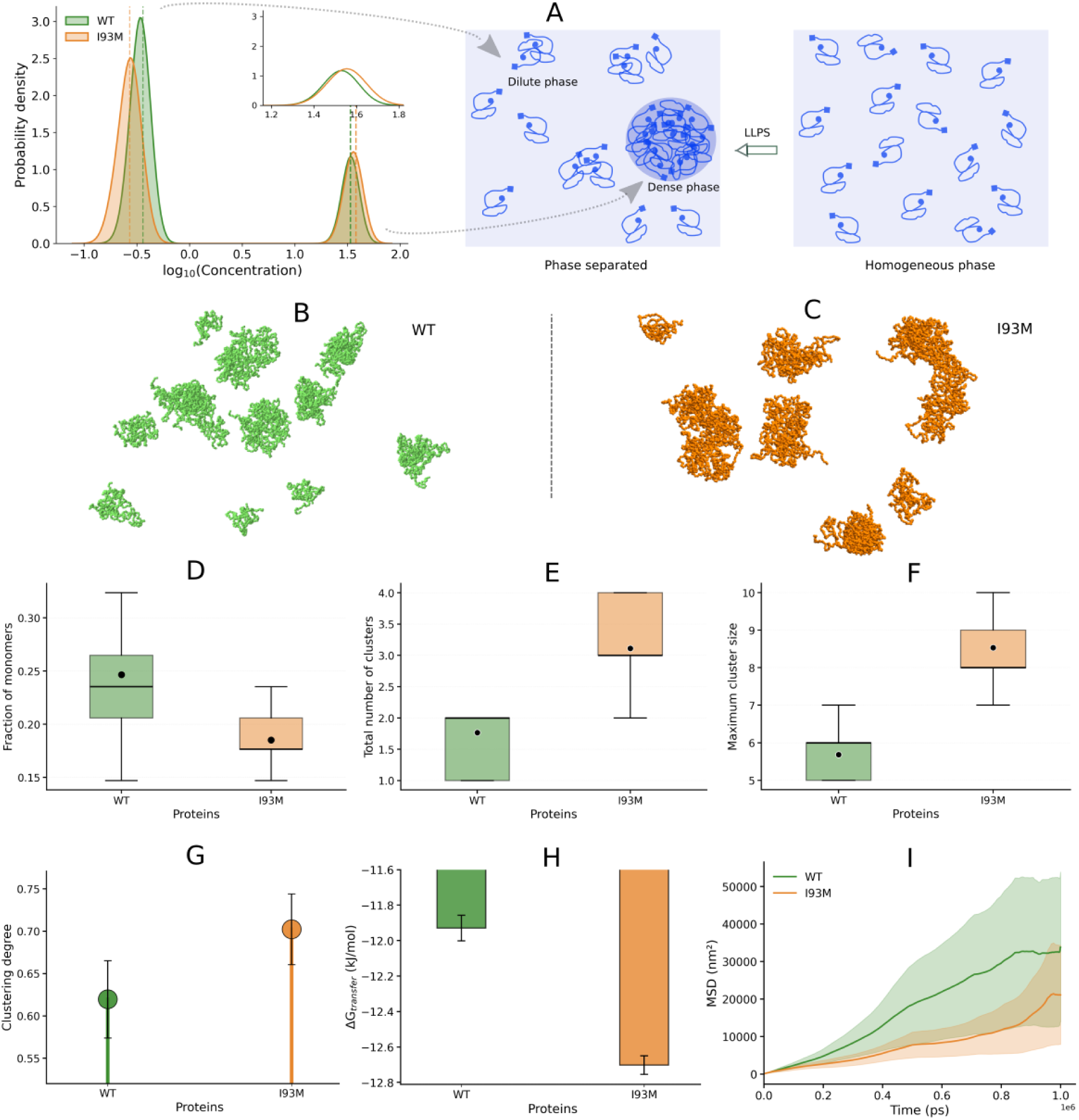
Phase separation behavior of WT and I93M UCH-L1. (A) Schematic illustration of LLPS, showing the transition from a homogeneous distribution of protein chains to coexistence of dilute and dense phases (right), along with corresponding protein concentration profiles represented as probability densities of log_10_ concentration (left) obtained from multichain CG simulations for WT and I93M. (B–C) Representative snapshots of multichain CG simulations for WT (B) and I93M (C), shown at equivalent simulation time points. (D) Box plots showing the fraction of monomeric proteins (number of monomers normalized by the total number of chains) in WT and I93M systems. The horizontal bold line denotes the median, box limits represent the interquartile range (IQR), and whiskers indicate the 5–95% range. Black dots indicate mean values. (E) Box plots showing the distribution of the total number of clusters formed in WT and I93M systems. The statistical representation follows the same definition as in panel (D). (F) Box plots showing the distribution of the maximum cluster size (number of chains in the largest cluster) in WT and I93M systems. The statistical representation follows the same definition as in panel (D). (G) Bar plot of clustering degree quantifying the extent of intermolecular self-assembly in WT and I93M systems. (H) ΔG*_trans_* associated with phase separation for WT and I93M. (I) Mean-squared displacement (MSD) of proteins in WT and I93M systems.

Quantitative analysis reinforces this conclusion across multiple descriptors. The I93M system exhibits a significantly lower fraction of monomeric proteins (Fig. 4D), a greater number of clusters (Fig. 4E), and larger maximum cluster sizes (Fig. 4F), indicative of more frequent oligomerization and larger condensate formation, demonstrating enhanced self-assembly and more extensive phase separation. The clustering degree[51]—defined as the ratio of chains in the dense phase (*N*_dense_) to total chains (*N*_total_) in system (Eq. 1)—is significantly higher for I93M (Fig. 4G), indicating a larger proportion of the total protein population incorporated into condensates.

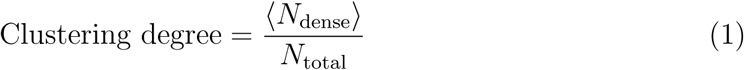

To evaluate the thermodynamic favorability of phase separation, we calculated the free energy of transfer (Δ*G*_trans_) for both systems[37], defined as the free energy cost for a protein chain to partition from the dilute into the dense phase computed from the concentrations of protein in the dilute (c_dilute_) and dense (c_dense_) phases (Eq. (2)):

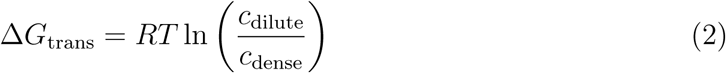

As shown in Fig. 4H, Δ*G*_trans_ is negative for both WT and I93M, indicating that partitioning of protein chains into the dense phase is thermodynamically favored in both cases. Notably, Δ*G*_trans_ is more negative for the I93M mutant, reflecting an increased thermodynamic driving force for transfer from the dilute to the dense phase relative to WT. Finally, mean-squared displacement (MSD) analysis reveals consistently lower values in the I93M system (Fig. 4I), indicating reduced protein mobility consistent with enhanced self-association and more extensive condensate formation. Together, these results demonstrate that the I93M mutation enhances LLPS across structural, thermodynamic, and kinetic descriptors.

### Molecular determinants of altered phase behavior

To identify the molecular basis of enhanced LLPS in the I93M mutant, we next examined intermolecular interaction patterns and intrachain organization in multichain simulations. The I93M mutation significantly increases the number of intermolecular contacts (Fig. 5A), indicating an enhanced propensity for interchain association. Consistently, difference maps of interresidue interchain contact probabilities (WT - I93M) reveal a widespread redistribution of intermolecular interaction patterns across the protein surface (Fig. 5B), with multiple regions exhibiting increased interchain contacts in the mutant. This suggests an overall shift toward enhanced intermolecular interaction propensity in the mutant ensemble. Energetic decomposition reveals that the enhanced phase separation is primarily driven by more favorable hydrophobic interactions, which are significantly more negative in the I93M system compared to WT, while Coulombic contributions remain largely unchanged (Fig. 5C). Consistently, solvation free-energy changes (ΔΔ*G*_solv_) shows a significantly reduced solvation penalty upon phase separation in the I93M mutant relative to WT (Fig. 5D), indicating thermodynamic stabilization of the condensed state.

**Figure 5:**
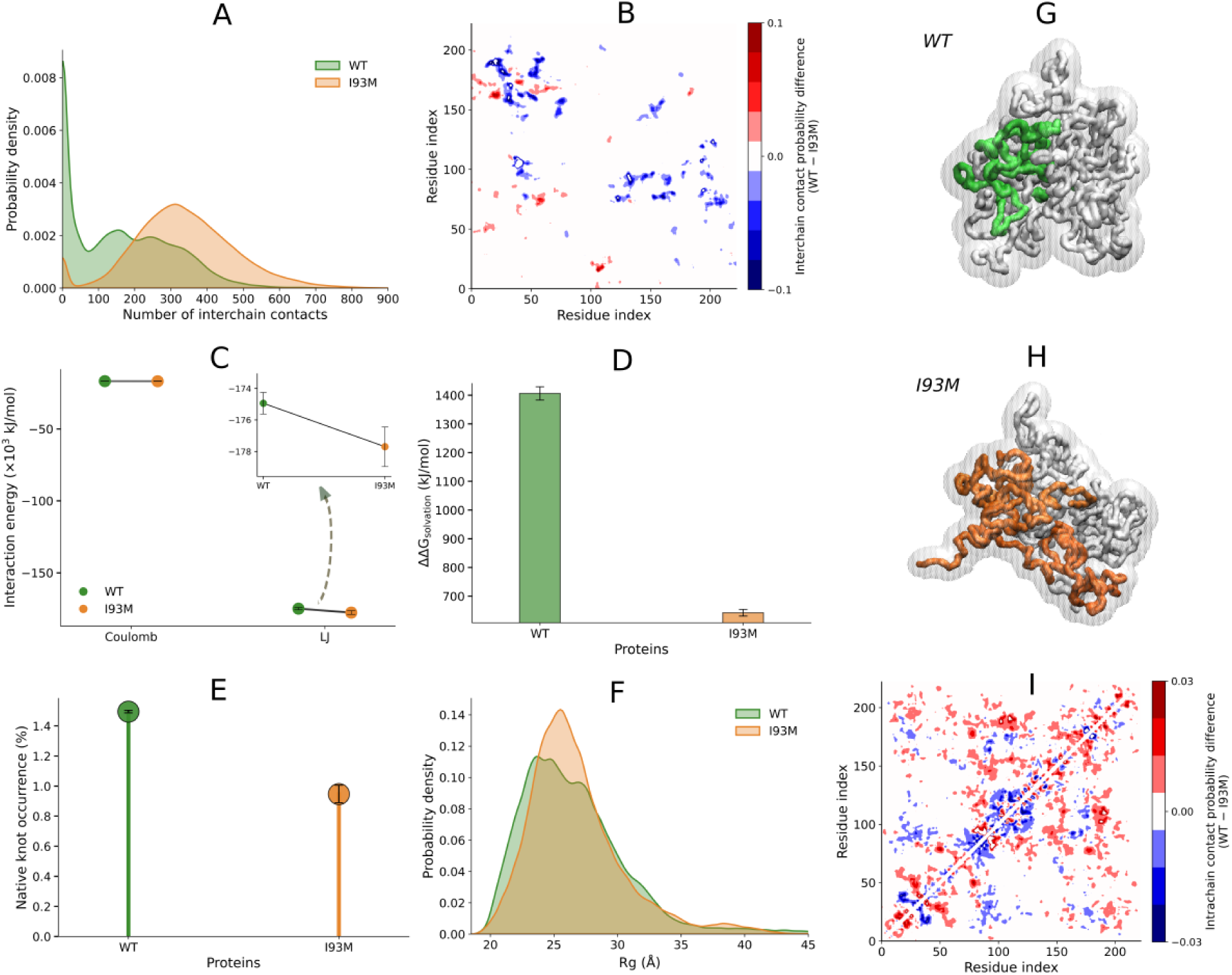
Structural and intermolecular determinants of altered phase behavior in I93M UCH-L1. (A) Distributions of the total number of interchain contacts for WT and I93M systems. (B) Difference maps of interresidue interchain contact probabilities (WT - I93M). The color scale (blue to red) indicates decreased (blue) and increased (red) contact probabilities in WT relative to I93M. (C) Energetic decomposition of protein interactions showing electrostatic and hydrophobic contributions for WT and I93M. Inset shows a zoomed view of the hydrophobic component. (D) Change in solvation free energy associated with phase separation (ΔΔG*_solv_*) for WT and I93M. (E) Percentage of native knot occurrence in WT and I93M systems. (F) R*_g_* distributions of protein chains in the dense phase from multichain simulations for WT and I93M. Corresponding distributions for monomeric (non-associated) chains within the same simulations are provided in the Supplementary Information (Fig. S1A). (G-H) Representative condensate snapshots for WT (G) and I93M (H) systems. One protein chain is highlighted in color while surrounding chains are shown in white surface representation to illustrate differences in chain conformational organization within the condensed phase. (I) Difference maps of intrachain contact probabilities for protein chains in the dense phase from multichain simulations (WT - I93M). The color scale (blue to red) indicates decreased (blue) and increased (red) contact probabilities in WT relative to I93M. Corresponding maps for monomeric chains within the same simulations are provided in the Supplementary Information (Fig. S1B).

Analysis of intrachain organization reveals the complementary side of this interaction redistribution. The I93M variant exhibits reduced probability of native knot occurrence (Fig. 5E) and increased R*_g_* values in the multichain context (Fig. 5F and Fig. S1A), consistent with partial destabilization of the native topological ensemble and relaxation of backbone constraints. Representative condensate snapshots further illustrate these differences in chain organization, with WT chains adopting comparatively compact conformations, whereas I93M chains remain more extended within the condensed phase (Fig. 5G-H). Difference maps of intrachain contact probabilities confirm that WT UCH-L1 exhibits stronger internal residue–residue interactions compared to the I93M mutant (Fig. 5I and Fig. S1B), consistent with greater stabilization of the native knotted ensemble in WT, while the I93M variant shows reduced intrachain contact probabilities indicating weakened intramolecular cohesion. Thus, the mutation shifts the balance between intrachain stabilization and intermolecular association: by weakening intramolecular contacts while strengthening interchain interactions, knot destabilization promotes a redistribution of interaction propensity toward multichain assembly, thereby favoring phase separation. This coupling between backbone topology and interaction energetics provides a mechanistic framework linking intramolecular organization to LLPS behavior and sets the stage for altered condensate-level properties.

### Altered Material Properties and Topological Reorganization in Mutant Condensates

Beyond differences in phase separation propensity, the I93M mutation leads to measurable changes in condensate organization and dynamics indicative of an altered material state[52, 53, 54]. The I93M mutant exhibits significantly reduced translational diffusion coefficients compared to the WT (Fig. 6A), indicating slower molecular motion and increased effective internal constraints within the condensed phase. In contrast, rotational relaxation remains relatively faster in I93M (Fig. 6B), suggesting a partial decoupling between translational and rotational dynamics consistent with viscoelastic-like behavior. Analysis of interchain contact lifetimes reveals a pronounced shift toward long-lived interchain interactions in I93M system (Fig. 6C), with a significant fraction of contacts persisting over extended timescales that would hinder molecular exchange and structural reorganization.

**Figure 6:**
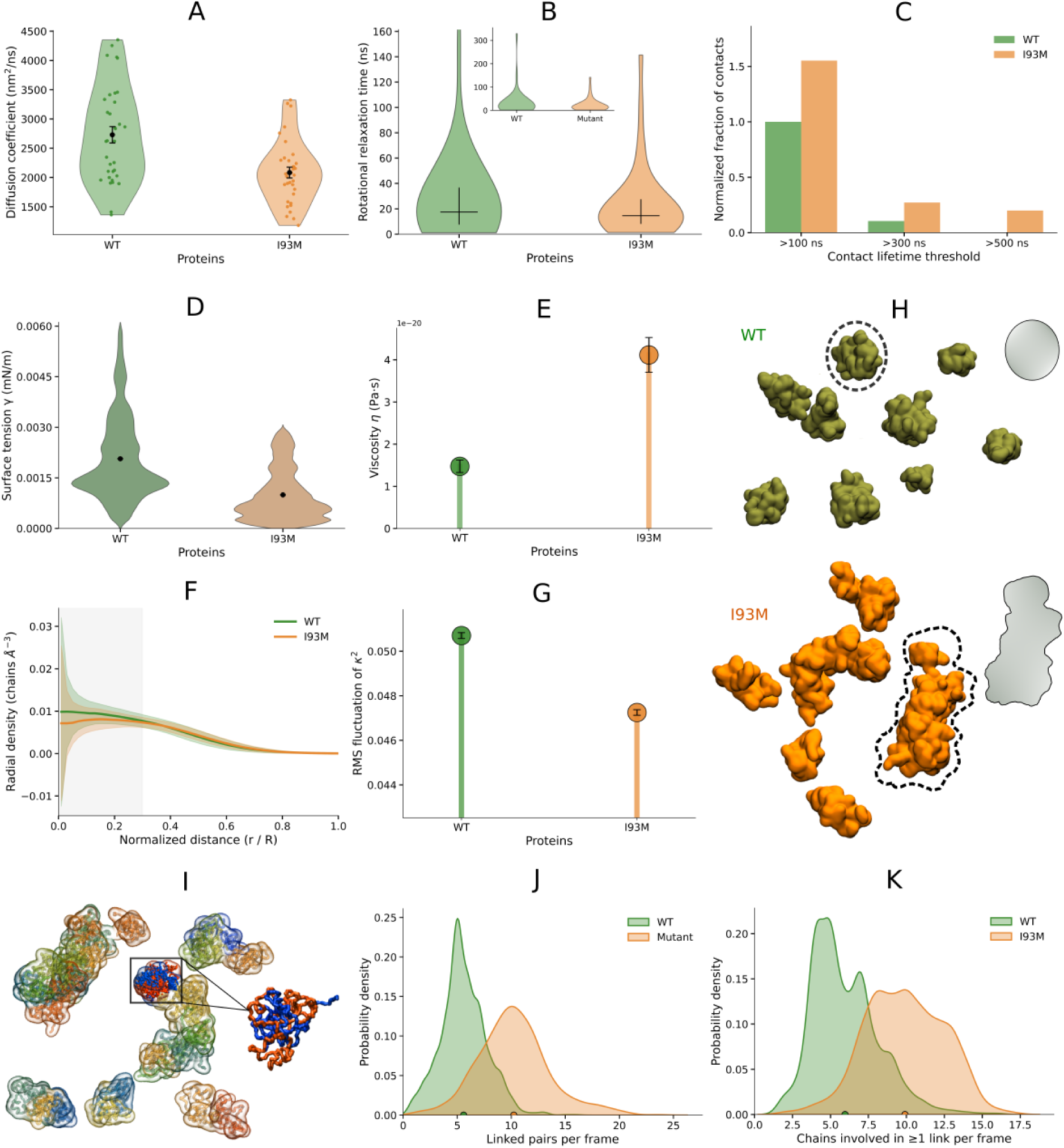
Altered material properties and topological remodeling in I93M condensates. (A) Violin plots of translational diffusion coefficients of proteins in condensates from multichain CG simulations for WT and I93M. (B) Violin plots of rotational relaxation times of proteins; inset shows a zoomed-out view. (C) Interchain contact lifetimes (D–E) Surface tension (Γ_20_; see Fig. S2 for Γ_22_) (D) and viscosity (E) of condensates, (F) Radial density profiles of condensates (G) Root-mean-square (RMS) shape fluctuations of condensates for WT and I93M. (H) Representative simulation snapshots of WT and I93M condensates, illustrating larger, asymmetric, and more gel-like assemblies in the mutant. (I) Snapshot highlighting interchain topological links in I93M condensates (see Gauss linking number analysis in Methods). (J) Probability distribution of the number of linked chain pairs in WT and I93M. (K) Probability distribution of the number of chains involved in one or more interchain links for WT and I93M.

These microscopic changes are reflected in emergent condensate material properties. The I93M variant exhibits reduced effective surface tension (Fig. 6D and Fig. S2), indicating weaker interfacial restoring forces, alongside markedly elevated effective viscosity (Fig. 6E), derived from shape anisotropy relaxation (see Methods), consistent with greater internal resistance to structural reorganization. Radial density profiles reveal a lower central density in mutant condensates (Fig. 6F), pointing to heterogeneous internal packing organization. Root-mean-square shape fluctuations are strongly suppressed in the mutant(Fig. 6G), indicating reduced droplet deformability despite sustained internal molecular motion — consistent with the preserved rotational relaxation noted above and suggestive of a dynamic hierarchy in which local motions persist while large-scale fluctuations are arrested. Collectively, these properties signal a transition toward viscoelastic, gel-like condensate behavior, further illustrated by the irregular and less deformable I93M condensate morphologies in representative snapshots (Fig. 6H).

A particularly striking feature of mutant condensates is the emergence of extensive interchain topological linking (Fig. 6I). Quantified via the Gauss linking number (see Methods), the I93M system forms significantly more linked chain pairs (Fig. 6J) with a higher fraction of chains participating in interchain links (Fig. 6K), consistent with an increased level of topological entanglement. This is further supported by the higher number of link-containing clusters observed in the mutant (Fig. S3), indicating that topological connectivity is more widespread across condensate structures. Importantly, this enhancement in interchain linking occurs in parallel with a reduction in intrachain knotting, suggesting a redistribution of topological organization from intramolecular to intermolecular degrees of freedom. This redistribution—from intrachain knotting to interchain entanglement—provides a mechanistic basis for the observed kinetic trapping: interchain topological links act as long-lived crosslinks that stabilize the condensate network, hinder relaxation, and promote persistent, aggregation-prone states more akin to gel-like assemblies than of dynamic liquid droplets.

## Discussion

Taken together, our results establish a mechanistic framework linking backbone knot stability, LLPS propensity, and emergent condensate material properties[52, 53, 54] in a disease-relevant protein. In WT UCH-L1, the deeply knotted topology functions as a conformational constraint that stabilizes intrachain contacts, limits conformational expansion, and suppresses excessive intermolecular association, resulting in dynamic, reversible condensates consistent with liquid-like behavior[34, 36, 53]. The I93M mutation disrupts this intrinsic topological constraint by destabilizing the native knot, shifting the conformational ensemble toward expanded states that are sparsely populated in WT[55, 56]. Under multichain conditions, these expanded states act as nucleation-competent species[2], that promote intermolecular contacts and dense-phase formation through both thermodynamic (enhanced hydrophobic interactions, reduced solvation penalty) and kinetic (long-lived contacts, interchain entanglement) pathways.

A central and unexpected finding is the coupling between intrachain topological destabilization and interchain topological entanglement in condensates. The reduction in native knot occupancy in the I93M mutant is accompanied by an increase in interchain linking within condensates, suggesting that the topological constraints released at the single-chain level are redistributed to the intermolecular level within the dense phase. This redistribution has direct material consequences: interchain topological links function as persistent crosslinks that suppress structural relaxation and promote viscoelastic, gel-like condensate behavior[57, 52, 54]. This mechanism may be broadly relevant, as topological entanglement between chains in dense phases is a recognized source of kinetic arrest[58, 35] in polymer systems, and its emergence in a disease-associated protein mutant suggests a previously unrecognized route to pathological condensate solidification[39, 41].

These findings introduce the concept of topological proteostasis: the maintenance of non-trivial backbone topology acts as a safeguard against pathological phase transitions[10, 53, 11]. In this framework, the knotted architecture of UCH-L1 is not merely a structural curiosity but an active functional element that buffers the protein against aberrant intermolecular association. Disruption of this protection—whether through mutation, post-translational modification, or cellular stress—may represent a general failure mode for knotted proteins, which constitute a small yet evolutionarily conserved class within the human proteome. The strict conservation of the 5_2_ knot across UCH-family deubiquitinases over large phylogenetic distances is consistent with positive selection for this topological protective function, beyond any contribution to enzymatic activity.

Finally, these results establish protein backbone topology as a previously unrecognized regulatory dimension for biomolecular condensates, complementing sequence composition, disorder content, and interaction valency. Topology is absent from existing sequence- or disorder-based models of phase separation, yet our simulations demonstrate that it can dominate phase behavior in a folded protein through its control over conformational entropy and the balance between intra- and intermolecular interactions. Whether approaches that reinforce native knot integrity, such as small molecules, chaperones, or other means, could modulate condensate material properties in a therapeutically meaningful way remains an open and experimentally tractable question[59, 60]. The mechanistic framework established here provides a foundation for future studies aimed at probing this possibility and more broadly at understanding how topological features of folded proteins shape the emergent behavior of biomolecular condensates.

## Models and Methods

### AA MD Simulations

WT UCH-L1 coordinates were taken from PDB ID 2ETL and the I93M mutant from PDB ID 3IRT. All AA MD simulations used the CHARMM36m force field for proteins and ions together with the CHARMM-modified TIP3P water model[61]. Systems were solvated in rectangular boxes (minimum 2 nm protein-to-boundary distance), neutralized, and supplemented with Na^+^ and Cl^−^ ions to 150 mM. Periodic boundary conditions were applied in all directions. After steepest-descent energy minimization, system were equilibrated under NVT conditions for 1 ns followed by NPT for 5 ns at 300 K and 1 bar. Temperature was controlled using the velocity-rescale thermostat, and pressure was maintained using the Parrinello–Rahman barostat. AA MD simulations are performed for each monomers and dimers systems (starting from pre-formed dimer structures) with both WT and I93M variant individually. In each case, production simulations were performed for at least 500 ns per replica using a 2 fs integration time step. Each of the simulations are replicated at least six times to maintain statistical reproducibility. All the simulations are performed with Gromacs software[62].

### CG MD Simulations

CG simulations used the Martini 3[63] force field. All CG systems were solvated in Martini water with 150 mM NaCl under periodic boundary conditions, energy-minimized using steepest descent, and equilibrated through sequential NVT using the velocity-rescale thermostat (5 ns) and NPT (5 ns) stages using a velocity-rescale thermostat coupled with a Berendsen barostat. Production simulations were performed using a velocity-Verlet integrator with a 0.02 ps time step, maintaining temperature and pressure with a velocity-rescale thermostat and Berendsen barostat, respectively. For CG monomer simulations of WT and I93M UCH-L1, each protein was placed in a cubic simulation box with a minimum distance of 2 nm from the protein to the box boundaries. For CG dimer simulations, two monomers were placed in a cubic box with initial COM separations exceeding the nonbonded interaction cutoff to preclude artificial initial contacts, and simulated to characterize encounter-based association kinetics. For each case production simulations were performed for at least 1 µs per replica and total six replica simulations are carried out to ensure statistical robustness. Multichain phase separation simulations used 34 protein copies randomly dispersed in a 48 × 48 × 48 nm^3^ box (∼500 µM protein concentration). Each system was simulated in three independent replicas for at least 2.5 µs, and analyses were performed on the final 1 µs of concatenated trajectories. All CG simulations were performed in GROMACS using the similar simulation protocol.

### Cluster Identification and Phase Separation Quantification

Protein clusters were identified using a distance-based clustering algorithm, where two chains were considered connected if any pair of backbone beads approached within 0.8 nm[37, 64]. Clusters ≥5 chains were classified as the dense phase; smaller assemblies were assigned to the dilute phase.

Protein concentration in each phase was calculated as[65]

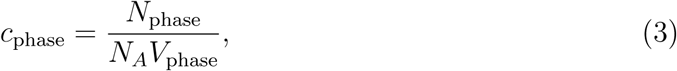

where *N*_phase_ is the number of protein chains in the phase, *N_A_*is Avogadro’s number, and *V*_phase_ is the volume occupied by that phase. Dense-phase volume was estimated from the eigenvalues *λ*_1_, *λ*_2_, and *λ*_3_ of the gyration tensor:

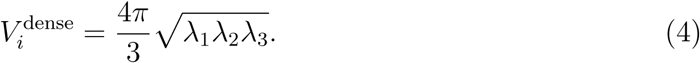

The dilute-phase volume was the remaining simulation box volume after subtracting all dense-droplet volumes.

### Knot Detection and Topological Analysis

Knot type was assigned frame-frame using the Jones polynomial invariant as implemented in Topoly [50, 66], with open backbone curves closed via a mass-centre scheme prior to invariant evaluation. The knotted core was identified by iterative terminal trimming to the minimal topology-preserving backbone segment; knot size was quantified as the number of C*α* atoms within this segment and knot position as the midpoint C*α* between the knot termini. Populations of knotted and unknotted states were aggregated across all replicas.

Inter-chain topological linking within dense-phase clusters was quantified using the Gauss linking number *L_k_*:

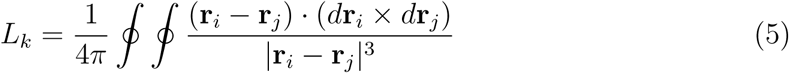

following standard formulations for polymer topology[67, 48], where **r***_i_* and **r***_j_* denote the positions along chains *i* and *j*, each closed via end-to-end closure. Chain pairs were classified as topologically linked when |*L_k_*| ≥ 0.5.

### Condensate Material Properties

Condensate shape anisotropy *κ*^2^ was computed from the eigenvalues *λ_i_* of the gyration tensor as:

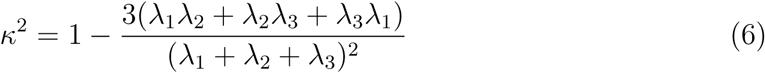

Relaxation times *τ* were obtained by fitting a single-exponential decay to the autocorrelation function of *κ*^2^ fluctuations. Effective viscosity *η* was estimated as:

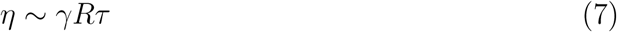

where *γ* is the surface tension and *R* is the droplet radius obtained from the radius of gyration of the dense phase, following Rayleigh–Lamb theory for overdamped interfacial relaxation [68, 69].

Surface tension was estimated from thermally driven shape fluctuations using a capillary fluctuation framework. Deviations of ellipsoidal semi-axes a and b from an equivalent-volume sphere of radius R were defined as:

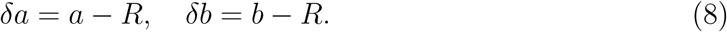

Assuming small fluctuations around a spherical equilibrium shape, the surface tension *γ* is given by:

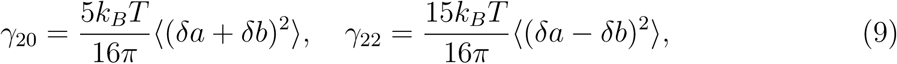

where where k*_B_* is the Boltzmann constant, T is the temperature, and angular brackets denote time averaging. The radius R was computed from the instantaneous droplet volume, and a,b were obtained from the eigenvalues of the gyration tensor of the dense phase. For isotropic droplets, *γ*_20_ and *γ*_22_ converge, and the surface tension was taken as *γ* ≈ *γ*_20_ ≈ *γ*_22_.

Interchain contacts were defined when backbone beads from distinct chains approached within 8 Å. Contact matrices and lifetimes were computed across trajectories to quantify network persistence. Contact lifetimes were classified as short (*<* 100 ns), intermediate (100–200 ns), or long-lived (*>* 500 ns).

### Software and Data Availability

All simulations were performed in GROMACS[62]. Analysis used GROMACS tools, MDAnalysis, NumPy, SciPy, and custom Python scripts. Analysis scripts are available upon reasonable request or via a public repository. All data supporting the findings are available within the Article and Supplementary Information.

## Acknowledgements

This work was supported in part by the National Institutes of Health through grant 5R01GM093290 and the U.S. Army Research Laboratory under the Cooperative Agreement No. W911NF-21-2-0007. Calculations were carried out using facilities created with the support of the U.S. National Science Foundation via Major Research Infrastructure grant numbers: 1625061 and 2216289.

## References

[1] Christopher A Ross and Michelle A Poirier. Protein aggregation and neurodegenerative disease. Nature medicine, 10(Suppl 7):S10–S17, 2004.

[2] Fabrizio Chiti and Christopher M Dobson. Protein misfolding, amyloid formation, and human disease: a summary of progress over the last decade. Annual review of biochemistry, 86:27–68, 2017.

[3] Tuomas PJ Knowles, Michele Vendruscolo, and Christopher M Dobson. The amyloid state and its association with protein misfolding diseases. Nature reviews Molecular cell biology, 15(6):384–396, 2014.

[4] Aaron Ciechanover and Yong Tae Kwon. Degradation of misfolded proteins in neurodegenerative diseases: therapeutic targets and strategies. Experimental & molecular medicine, 47(3):e147–e147, 2015.

[5] David Komander, Michael J Clague, and Sylvie Urbé. Breaking the chains: structure and function of the deubiquitinases. Nature reviews Molecular cell biology, 10(8): 550–563, 2009.

[6] Julia Reichelt, Wiebke Sachs, Sarah Frömbling, Julia Fehlert, Maja Studencka-Turski, Anna Betz, Desiree Loreth, Lukas Blume, Susanne Witt, Sandra Pohl, et al. Non-functional ubiquitin c-terminal hydrolase l1 drives podocyte injury through impairing proteasomes in autoimmune glomerulonephritis. Nature communications, 14 (1):2114, 2023.

[7] Christian Grethe, Mirko Schmidt, Gian-Marvin Kipka, Rachel O’Dea, Kai Gallant, Petra Janning, and Malte Gersch. Structural basis for specific inhibition of the deubiquitinase uchl1. Nature Communications, 13(1):5950, 2022.

[8] Yoko Goto, Lihua Zeng, Chan Joo Yeom, Yuxi Zhu, Akiyo Morinibu, Kazumi Shinomiya, Minoru Kobayashi, Kiichi Hirota, Satoshi Itasaka, Michio Yoshimura, et al. Uchl1 provides diagnostic and antimetastatic strategies due to its deubiquitinating effect on hif-1*α*. Nature communications, 6(1):6153, 2015.

[9] Zhihua Liu, Robin K Meray, Tom N Grammatopoulos, Ross A Fredenburg, Mark R Cookson, Yichin Liu, Todd Logan, and Peter T Lansbury Jr. Membrane-associated farnesylated uch-l1 promotes *α*-synuclein neurotoxicity and is a therapeutic target for parkinson’s disease. Proceedings of the National Academy of Sciences, 106(12): 4635–4640, 2009.

[10] William E Balch, Richard I Morimoto, Andrew Dillin, and Jeffery W Kelly. Adapting proteostasis for disease intervention. science, 319(5865):916–919, 2008.

[11] Susmita Kaushik and Ana Maria Cuervo. Proteostasis and aging. Nature medicine, 21(12):1406–1415, 2015.

[12] Keith D Wilkinson, Keunmyoung Lee, Seema Deshpande, Penelope Duerksen-Hughes, Jeremy M Boss, and Jan Pohl. The neuron-specific protein pgp 9.5 is a ubiquitin carboxyl-terminal hydrolase. Science, 246(4930):670–673,1989.

[13] Kaya Bilguvar, Navneet K Tyagi, Cigdem Ozkara, Beyhan Tuysuz, Mehmet Bakircioglu, Murim Choi, Sakir Delil, Ahmet O Caglayan, Jacob F Baranoski, Ozdem Erturk, et al. Recessive loss of function of the neuronal ubiquitin hydrolase uchl1 leads to early-onset progressive neurodegeneration. Proceedings of the National Academy of Sciences, 110(9):3489–3494, 2013.

[14] Chittaranjan Das, Quyen Q Hoang, Cheryl A Kreinbring, Sarah J Luchansky, Robin K Meray, Soumya S Ray, Peter T Lansbury, Dagmar Ringe, and Gregory A Petsko. Structural basis for conformational plasticity of the parkinson’s disease-associated ubiquitin hydrolase uch-l1. Proceedings of the National Academy of Sciences, 103(12):4675–4680, 2006.

[15] Fujun Chen, Yoshie Sugiura, Kalisa Galina Myers, Yun Liu, and Weichun Lin. Ubiquitin carboxyl-terminal hydrolase l1 is required for maintaining the structure and function of the neuromuscular junction. Proceedings of the National Academy of Sciences, 107(4):1636–1641, 2010.

[16] Daewon Lee, Eunju Yoon, Su Jin Ham, Kunwoo Lee, Hansaem Jang, Daihn Woo, Da Hyun Lee, Sehyeon Kim, Sekyu Choi, and Jongkyeong Chung. Diabetic sensory neuropathy and insulin resistance are induced by loss of uchl1 in drosophila. Nature Communications, 15(1):468, 2024.

[17] Anna T Reinicke, Karoline Laban, Marlies Sachs, Vanessa Kraus, Michael Walden, Markus Damme, Wiebke Sachs, Julia Reichelt, Michaela Schweizer, Philipp Christoph Janiesch, et al. Ubiquitin c-terminal hydrolase l1 (uch-l1) loss causes neurodegeneration by altering protein turnover in the first postnatal weeks. Proceedings of the National Academy of Sciences, 116(16):7963–7972, 2019.

[18] Hao Liu, Nadya Povysheva, Marie E Rose, Zhiping Mi, Joseph S Banton, Wenjin Li, Fenghua Chen, Daniel P Reay, Germán Barrionuevo, Feng Zhang, et al. Role of uchl1 in axonal injury and functional recovery after cerebral ischemia. Proceedings of the National Academy of Sciences, 116(10):4643–4650, 2019.

[19] Leonardus MI Koharudin, Hao Liu, Roberto Di Maio, Ravindra B Kodali, Steven H Graham, and Angela M Gronenborn. Cyclopentenone prostaglandin-induced unfolding and aggregation of the parkinson disease-associated uch-l1. Proceedings of the National Academy of Sciences, 107(15):6835–6840, 2010.

[20] Elisabeth Leroy, Rebecca Boyer, Georg Auburger, Barbara Leube, Gudrun Ulm, Eva Mezey, Gyongyi Harta, Michael J. Brownstein, Sobhanadditya Jonnalagada, Tanya Chernova, Anindya Dehejia, Christian Lavedan, Thomas Gasser, Peter J. Steinbach, Keith D. Wilkinson, and Mihael H. Polymeropoulos. The ubiquitin pathway in parkinson’s disease. Nature, 395(6701):451–452, 1998.

[21] Fabian Ziegler, Nicole CH Lim, Soumit Sankar Mandal, Benjamin Pelz, Wei-Ping Ng, Michael Schlierf, Sophie E Jackson, and Matthias Rief. Knotting and unknotting of a protein in single molecule experiments. Proceedings of the National Academy of Sciences, 113(27):7533–7538, 2016.

[22] Paul Bishop, Dan Rocca, and Jeremy M Henley. Ubiquitin c-terminal hydrolase l1 (uch-l1): structure, distribution and roles in brain function and dysfunction. Biochemical Journal, 473(16):2453–2462, 2016.

[23] Susmita Sarkar, Mark DelloStritto, and Michael L Klein. Temperature-driven two-stage knot dynamics. Macromolecules, 2026.

[24] Sara GF Ferreira, Manoj K Sriramoju, Shang-Te Danny Hsu, Patricia FN Faisca, and Miguel Machuqueiro. Is there a functional role for the knotted topology in protein uch-l1? Journal of Chemical Information and Modeling, 64(17):6827–6837, 2024.

[25] William R Taylor. A deeply knotted protein structure and how it might fold. Nature, 406(6798):916–919, 2000.

[26] Chengzhi Liang and Kurt Mislow. Knots in proteins. Journal of the American Chemical Society, 116(24):11189–11190, 1994.

[27] Marc L Mansfield. Are there knots in proteins? Nature structural biology, 1(4): 213–214, 1994.

[28] Pawel Dabrowski-Tumanski and Joanna I Sulkowska. Topological knots and links in proteins. Proceedings of the National Academy of Sciences, 114(13):3415–3420, 2017.

[29] Rhonald C Lua and Alexander Y Grosberg. Statistics of knots, geometry of conformations, and evolution of proteins. PLoS computational biology, 2(5):e45, 2006.

[30] Peter Virnau, Yacov Kantor, and Mehran Kardar. Knots in globule and coil phases of a model polyethylene. Journal of the American Chemical Society, 127(43):15102–15106, 2005.

[31] Wenquan Liang, Ru Feng, Xiaojia Li, Xingwei Duan, Shourui Feng, Jun Chen, Yicheng Li, Junqi Chen, Zezheng Liu, Xiaogang Wang, et al. A rankl-uchl1-scd13 negative feedback loop limits osteoclastogenesis in subchondral bone to prevent osteoarthritis progression. Nature communications, 15(1):8792, 2024.

[32] Alex Kluber, Timothy A Burt, and Cecilia Clementi. Size and topology modulate the effects of frustration in protein folding. Proceedings of the National Academy of Sciences, 115(37):9234–9239, 2018.

[33] Franco O Tzul, Daniel Vasilchuk, and George I Makhatadze. Evidence for the principle of minimal frustration in the evolution of protein folding landscapes. Proceedings of the National Academy of Sciences, 114(9):E1627–E1632, 2017.

[34] Clifford P Brangwynne, Christian R Eckmann, David S Courson, Agata Rybarska, Carsten Hoege, Jöbin Gharakhani, Frank Jülicher, and Anthony A Hyman. Germline p granules are liquid droplets that localize by controlled dissolution/condensation. Science, 324(5935):1729–1732, 2009.

[35] Salman F Banani, Hyun O Lee, Anthony A Hyman, and Michael K Rosen. Biomolecular condensates: organizers of cellular biochemistry. Nature reviews Molecular cell biology, 18(5):285–298, 2017.

[36] Yongdae Shin and Clifford P Brangwynne. Liquid phase condensation in cell physiology and disease. Science, 357(6357):eaaf4382, 2017.

[37] Susmita Sarkar and Jagannath Mondal. How salt and temperature drive reentrant condensation of a*β*40. Biochemistry, 63(22):3030–3044, 2024.

[38] Susmita Sarkar, Tharangattu N Narayanan, and Jagannath Mondal. A synergistic view on osmolyte’s role against salt and cold stress in biointerfaces. Langmuir, 39 (49):17581–17592, 2023.

[39] Avinash Patel, Hyun O Lee, Louise Jawerth, Shovamayee Maharana, Marcus Jahnel, Marco Y Hein, Stoyno Stoynov, Julia Mahamid, Shambaditya Saha, Titus M Franzmann, et al. A liquid-to-solid phase transition of the als protein fus accelerated by disease mutation. Cell, 162(5):1066–1077, 2015.

[40] Susmita Sarkar, Anku Guha, Rayantan Sadhukhan, Tharangattu N Narayanan, and Jagannath Mondal. Osmolytes as cryoprotectants under salt stress. ACS Biomaterials Science & Engineering, 9(10):5639–5652, 2023.

[41] Amandine Molliex, Jamshid Temirov, Jihun Lee, Maura Coughlin, Anderson P Kanagaraj, Hong Joo Kim, Tanja Mittag, and J Paul Taylor. Phase separation by low complexity domains promotes stress granule assembly and drives pathological fibrillization. Cell, 163(1):123–133, 2015.

[42] Susmita Sarkar, Rayantan Sadhukhan, Nandita Mohandas, Amogh K Ravi, Tharangattu N Narayanan, and Jagannath Mondal. Adenosine triphosphate inhibits cold-responsive aggregation. Langmuir, 40(41):21587–21599, 2024.

[43] Erik W Martin, Alex S Holehouse, Christy R Grace, Alex Hughes, Rohit V Pappu, and Tanja Mittag. Sequence determinants of the conformational properties of an intrinsically disordered protein prior to and upon multisite phosphorylation. Journal of the American Chemical Society, 138(47):15323–15335, 2016.

[44] Zhitao Liao, Bowen Jia, Dongshi Guan, Xudong Chen, Mingjie Zhang, and Penger Tong. Emergent mechanics of a networked multivalent protein condensate. Nature Communications, 16(1):5237, 2025.

[45] Susmita Sarkar, Anku Guha, Tharangattu N Narayanan, and Jagannath Mondal. Zwitterionic osmolytes revive surface charges under salt stress via dual mechanisms. The Journal of Physical Chemistry Letters, 13(24):5660–5668, 2022.

[46] Erik W Martin, Alex S Holehouse, Ivan Peran, Mina Farag, J Jeremias Incicco, Anne Bremer, Christy R Grace, Andrea Soranno, Rohit V Pappu, and Tanja Mittag. Valence and patterning of aromatic residues determine the phase behavior of prion-like domains. Science, 367(6478):694–699, 2020.

[47] Susmita Sarkar, Anku Guha, Tharangattu N Narayanan, and Jagannath Mondal. Osmolyte-induced modulation of hofmeister series. The Journal of Physical Chemistry B, 128(39):9436–9446, 2024.

[48] Cristian Micheletti, Davide Marenduzzo, and Enzo Orlandini. Polymers with spatial or topological constraints: Theoretical and computational results. Physics Reports, 504(1):1–73, 2011.

[49] Maziar Heidari, Helmut Schiessel, and Alireza Mashaghi. Circuit topology analysis of polymer folding reactions. ACS Central Science, 6(6):839–847, 2020.

[50] Kasturi Barkataki and Eleni Panagiotou. The jones polynomial of collections of open curves in 3-space. Proceedings of the Royal Society A: Mathematical, Physical and Engineering Sciences, 478(2267), 2022.

[51] Yiming Tang, Santu Bera, Yifei Yao, Jiyuan Zeng, Zenghui Lao, Xuewei Dong, Ehud Gazit, and Guanghong Wei. Prediction and characterization of liquid-liquid phase separation of minimalistic peptides. Cell Reports Physical Science, 2(9), 2021.

[52] Huabin Zhou, Jan Huertas, M Julia Maristany, Kieran Russell, June Ho Hwang, Run-Wen Yao, Nirnay Samanta, Joshua Hutchings, Ramya Billur, Momoko Shiozaki, et al. Multiscale structure of chromatin condensates explains phase separation and material properties. Science, 390(6777):eadv6588, 2025.

[53] Simon Alberti and Anthony A Hyman. Biomolecular condensates at the nexus of cellular stress, protein aggregation disease and ageing. Nature reviews Molecular cell biology, 22(3):196–213, 2021.

[54] Louise Jawerth, Elisabeth Fischer-Friedrich, Suropriya Saha, Jie Wang, Titus Franzmann, Xiaojie Zhang, Jenny Sachweh, Martine Ruer, Mahdiye Ijavi, Shambaditya Saha, et al. Protein condensates as aging maxwell fluids. Science, 370(6522):1317–1323, 2020.

[55] Andrew J Baldwin and Lewis E Kay. Nmr spectroscopy brings invisible protein states into focus. Nature chemical biology, 5(11):808–814, 2009.

[56] Philipp Neudecker, Paul Robustelli, Andrea Cavalli, Patrick Walsh, Patrik Lund-ström, Arash Zarrine-Afsar, Simon Sharpe, Michele Vendruscolo, and Lewis E Kay. Structure of an intermediate state in protein folding and aggregation. Science, 336 (6079):362–366, 2012.

[57] Clifford P Brangwynne, Peter Tompa, and Rohit V Pappu. Polymer physics of intracellular phase transitions. Nature Physics, 11(11):899–904, 2015.

[58] Jeong-Mo Choi, Alex S Holehouse, and Rohit V Pappu. Physical principles underlying the complex biology of intracellular phase transitions. Annual review of biophysics, 49(1):107–133, 2020.

[59] Prajwal Ciryam, Gian Gaetano Tartaglia, Richard I Morimoto, Christopher M Dobson, and Michele Vendruscolo. Widespread aggregation and neurodegenerative diseases are associated with supersaturated proteins. Cell reports, 5(3):781–790, 2013.

[60] Macy A. Seijo, Anisha Banerjee, and Caesar M. Hernandez. Network-level organization of systemic inflammation reflects early alzheimer’s-like behavioral changes. Scientific Reports, 16(1), 2025.

[61] Alex D MacKerell Jr, Donald Bashford, MLDR Bellott, Roland Leslie Dunbrack Jr, Jeffrey D Evanseck, Martin J Field, Stefan Fischer, Jiali Gao, Houyang Guo, Sookhee Ha, et al. All-atom empirical potential for molecular modeling and dynamics studies of proteins. The journal of physical chemistry B, 102(18):3586–3616, 1998.

[62] Mark James AbrI am preI aaham, Teemu Murtola, Roland Schulz, Szilárd Páll, Jeremy C Smith, Berk Hess, and Erik Lindahl. Gromacs: High performance molecular simulations through multi-level parallelism from laptops to supercomputers. SoftwareX, 1:19–25, 2015.

[63] Paulo CT Souza, Riccardo Alessandri, Jonathan Barnoud, Sebastian Thallmair, Ignacio Faustino, Fabian Grünewald, Ilias Patmanidis, Haleh Abdizadeh, Bart MH Bruininks, Tsjerk A Wassenaar, et al. Martini 3: a general purpose force field for coarse-grained molecular dynamics. Nature methods, 18(4):382–388, 2021.

[64] Susmita Sarkar, Saurabh Gupta, Chiranjit Mahato, Dibyendu Das, and Jagannath Mondal. Elucidating atp’s role as solubilizer of biomolecular aggregate. eLife, pages 13, RP99150, 2024.

[65] Hung T Nguyen, Naoto Hori, and D Thirumalai. Condensates in rna repeat sequences are heterogeneously organized and exhibit reptation dynamics. Nature chemistry, 14 (7):775–785, 2022.

[66] Pawel Dabrowski-Tumanski, Pawel Rubach, Wanda Niemyska, Bartosz Ambrozy Gren, and Joanna Ida Sulkowska. Topoly: Python package to analyze topology of polymers. Briefings in Bioinformatics, 22(3):bbaa196, 2021. doi: 10.1093/bib/bbaa196.

[67] Konstantin Klenin and Jörg Langowski. Computation of writhe in modeling of supercoiled dna. Biopolymers: Original Research on Biomolecules, 54(5):307–317, 2000.

[68] John Shipley Rowlinson and Benjamin Widom. Molecular theory of capillarity. Courier Corporation, 2013.

[69] Howard A Stone and L Gary Leal. The effects of surfactants on drop deformation and breakup. Journal of Fluid Mechanics, 220:161–186, 1990.

