## Supporting information for "Tightly Knotted Enzymes Inhibit Protein-Protein Aggregation"

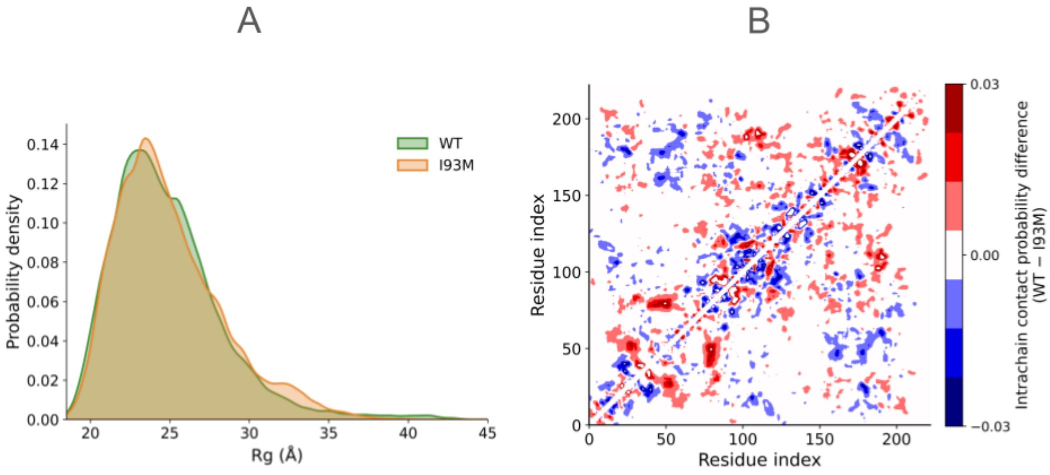


Figure S1: (A) $R_g$ distributions of protein chains in the monomeric form in dilute phase from multichain simulations for WT and I93M. (B) Difference maps of intrachain contact probabilities between WT and I93M for protein chains in the monomeric form in dilute phase from multichain simulations, calculated as WT - I93M. The color scale (blue to red) indicates reduced (blue) and increased (red) contact probabilities in WT relative to I93M.

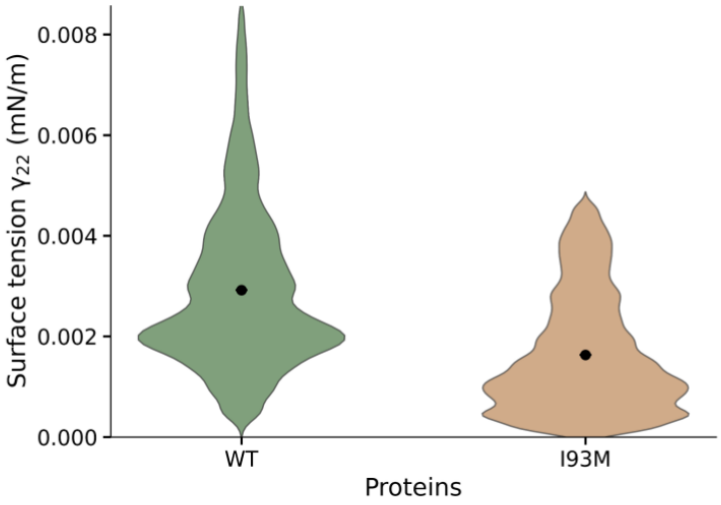


Figure S2: Surface tension ($\Gamma_{22}$) of condensates, indicating reduced mobility and increased resistance to flow in I93M.

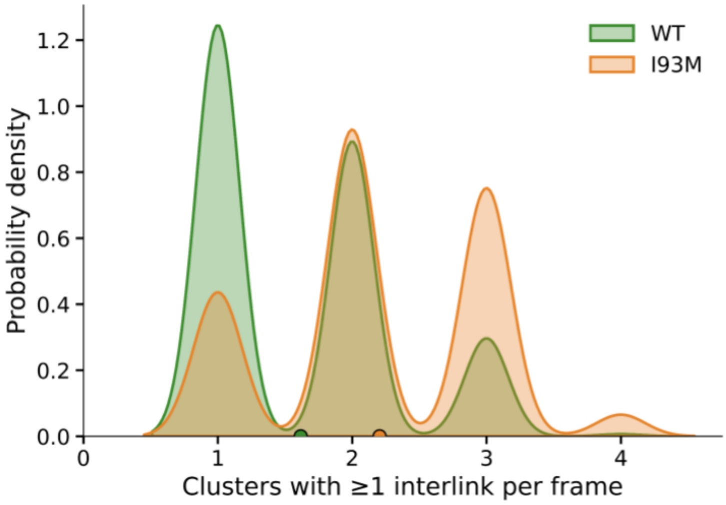


Figure S3: Probability distribution of the number of clusters with one or more than one interlinking per frame in WT and I93M.
